# Influence of residual microbial nucleic acid in instruments for hip joint arthroplasty on false-positive results of metagenomic sequencing

**DOI:** 10.1101/2022.03.25.485773

**Authors:** Han Yin, Chengtan Wang, Duliang Xu, Mingtao Xu, Chengzhi Ha, Wei Li, Feng Pang, Dawei Wang

**Author notes:** These authors contributed equally to this work (Han Yin, Chengtan Wang).

## Abstract

The purpose of this study was to analyze the effect of residual microbial nucleic acid in instruments for hip joint arthroplasty on false-positive sequencing results. Samples were taken from 3 different acetabular reamer for hip arthroplasty in 7 different hospitals. The whole process was strictly aseptic, metagenomic next-generation sequencing (mNGS) was performed according to standard operating procedures. The sterility of instruments was confirmed by culture method. The sequencing results of specimens from different hospitals were compared to analyze the difference of background bacteria. Bioinformatics analysis and visualization were presented through R language. A total of 26 samples were processed by mNGS, including 24 instrument swab samples, 1 blank swab control, and 1 blank water control. 254,314,707 reads were sequenced in all samples. The results showed that 1.13% of Clean Reads can be matched to pathogenic microorganism genomes, of which bacterial sequences account for 87.48%, fungal sequences account for 11.18%, parasite sequences account for 1.26%, and virus sequences account for 0.06%. The results of PCA (Principal Component Analysis) demonstrated that the distribution of bacteria on the surface of instruments was significantly different between medical institutions. Through the Venn diagram, it was found that 465 species of bacteria in all region hospitals, Liaocheng People’s Hospital had a maximum of 340 species of bacteria, followed by Guanxian County People’s Hospital with 169 species. The clustering heat map illustrated that the distribution of bacterial groups in three different instrument samples in the same hospital was basically the same, and the bacterial genera varied significantly among hospitals. The residual microbial nucleic acid fragments are mainly bacterial DNA and represent differences in different medical institutions, The establishment of independent background bacterial libraries in different medical institutions can effectively improve the accuracy of mNGS diagnosis and help to exclude background microorganisms interference.

## Introduction

Periprosthetic joint infection (PJI) is one of the serious complications after total joint replacement <u>[1]</u>. Timely and accurate microbiological diagnosis is helpful for effective infection control <u>[2]</u>. False-negative results due to low activity or low concentrations of bacteria, and difficult-to-cultivate pathogens in vitro are common limitations of this traditional method <u>[3, 4, 5, 6, 7]</u>. The metagenomic next-generation sequencing (mNGS) is a molecular detection method for pathogens that have attracted great attention in recent years. This method detects all pathogenic nucleic acid fragments in a specimen by unbiased shotgun sequencing and prompts its correlation with clinical infection. The mNGS technology has demonstrated broad application in pathogen identification of PJI cases <u>[2, 8, 9, 10]</u>. Throughimproving the sensitivity of pathogen detection, mNGS has a positive impact on improving the prognosis of PJI patients, rational use of antibiotics, and reducing the economic burden of patients <u>[11]</u>.

Although mNGS plays an increasingly important role in the diagnosis of PJI, accurate interpretation of sequencing reports is a difficult task for clinicians and laboratory reporters. Due to the extremely high sensitivity of mNGS testing, even a small amount of nucleic acid fragments exposed during sampling or testing may lead to false positives, which is one of the biggest challenges in interpreting mNGS testing reports. Street *et al.* concluded that high levels of DNA contamination may be introduced in the process of mNGS, which is difficult to remove and can only be minimized during the operation <u>[12]</u>. Huang *et al.* also reported the interference of exogenous DNA contamination in the detection of bone and joint infections using mNGS <u>[13]</u>. Detecting contaminating nucleic acid sequences during mNGS and mistaking them for pathogenic microorganisms may be a common problem for all inspectors <u>[14]</u>.

In this study, for the first time, we experimentally detected and established PJI-related interfering nucleic acid background microbial libraries (BML) in different medical institutions. We obtained residual nucleic acid on the surface of instruments for joint arthroplastyin different medical institutions through mNGS, and applied bioinformatics methods to analyze and compare the differences in microbial distribution. To more accurately apply mNGS to diagnose PJI diseases, each medical institution must establish its BML database.

## Materials and methods

### Ethics Statement

This study does not involve human or animal samples. According to the “Measures for Ethical Review of Biomedical Research Involving Humans” of the People’s Republic of China, this study does not require ethical permission. All samples used in this study were anonymized. All experiments were performed according to the Declaration of Helsinki.

### Experimental Design and Sample Acquisition

Samples were taken from 7 hospitals in Liaocheng City (Shandong Province, China), including Liaocheng People’s Hospital (No. LPH 1-3, and No. LPH 4-6 containing human Hela cells), Liaocheng Hospital of Traditional Chinese Medicine (No. LHTCM 1-3), Dong’e County People’s Hospital (No. DCPH 1-3), Guanxian County People’s Hospital (No. GCPH 1-3), Yanggu County People’s Hospital (No. YCPH 1-3), Shenxian County People’s Hospital (No. SCPH 1-3), Chaocheng County People’s Hospital (No. CCPH 1-3). Three different acetabular reamer for hip arthroplasty were collected from each hospital. Among them, Liaocheng People’s Hospital took samples twice. In one experiment, Hela cells were added to determine the effect of adding human sequence (LPH4-6), and the other was directly sequenced (LPH1-3). The samples from the other 6 hospitals were directly sequenced without adding human sequences.

All specimens were collected in a thousand-level laminar flow operating room that had constant temperature and humidity to minimize contamination. Sampling personnel are dressed in accordance with sterile protection requirements, with hats, masks, and gloves. Throughout the procedure, avoid talking to prevent possible dental bacterial contamination. Before aseptic surgery, take a special swab to wipe the surface of the acetabular reamer, and mix it with sterile water after wiping. The treated liquid was divided into two parts as sample treatment solution, one for microbial culture and the other for sequencing. In addition to the conventional internal reference and quality control substances in the sequencing process, this experiment set up a swab (Control 1) and ultrapure water (Control 2) without wiping any instruments involved as additional control substances. All controls follow the normal test procedure until the report is obtained.

### Microbial Culture

The sample treatment solution was inoculated in 0.1 ml aliquots on aerobic blood agar, anaerobic blood agar, and fungal Sabouraud agar. Then they were incubated under the corresponding temperature and environmental conditions, respectively, 35°C and CO_2_ content of 5-7% for 6 days aerobic culture, 14 days in the anaerobic tank for anaerobic culture, and 28°C incubators for fungus medium 28 days. Additionally, 1 ml of the treatment solution was injected into the BACTEC Peds Plus/F bottle and incubated for 6 days in the BACTEC FX200 automated system. Furthermore, mycobacterial cultures were performed in liquid cultures for 42 days by a BACTEC MGIT 960 system (Becton-Dickinson, USA). Any growth from the treatment fluid was considered positive.

### mNGS Operation Process

The collected samples were subjected to mNGS of DNA by PMSEQ products at BGI PathoGenesis 50 (MGI Tech Co., Ltd, Wuhan, China). For complete information on sample preparation, sequencing methods, and metagenomic data analysis, see Supplementary Methods.

### Data Aggregation and Visualization

Bioinformatics analysis and visualization using the relative abundance of different microbial genera as indicators are all realized by R language. The principal component analysis uses the “PCA tools” R package; heatmaps use the “pheatmap” R package. Other analysis and data preprocessing use R language built-in functions or the “ggplot2” R package.

## Results

### Microbial Culture

Aerobic and anaerobic cultures of bacteria were negative after the designed number of culture days. Simultaneously, the enrichment operation in which the treatment solution was injected into the culture bottle did not obtain a positive culture result. Fungal and mycobacterial cultures were also negative.

### mNGS Sequencing Results

A total of 26 samples from 7 different medical institutions were included in this study, including 24 instrument swab samples and 1 blank control (Control 1), and 1 blank water control (Control 2). In these 26 samples, 254,314,707 fragment reads were obtained by sequencing, of which 25,210,926 were internal reference sequences (detection rate of internal reference was 100%). The Clean Reads obtained after removing adapter sequences and low-quality sequencing data were aligned to the human reference genome, finally, 88.31% could be aligned to the human reference genome, 10.56% were not aligned to the specific genome, and 1.13% compared to pathogenic microorganism genomes (Fig 1A), of which bacterial sequences account for 87.48%, fungal sequences account for 11.18%, parasite sequences account for 1.27%, and virus sequences account for 0.06% (Fig 1B).

**Fig 1.**
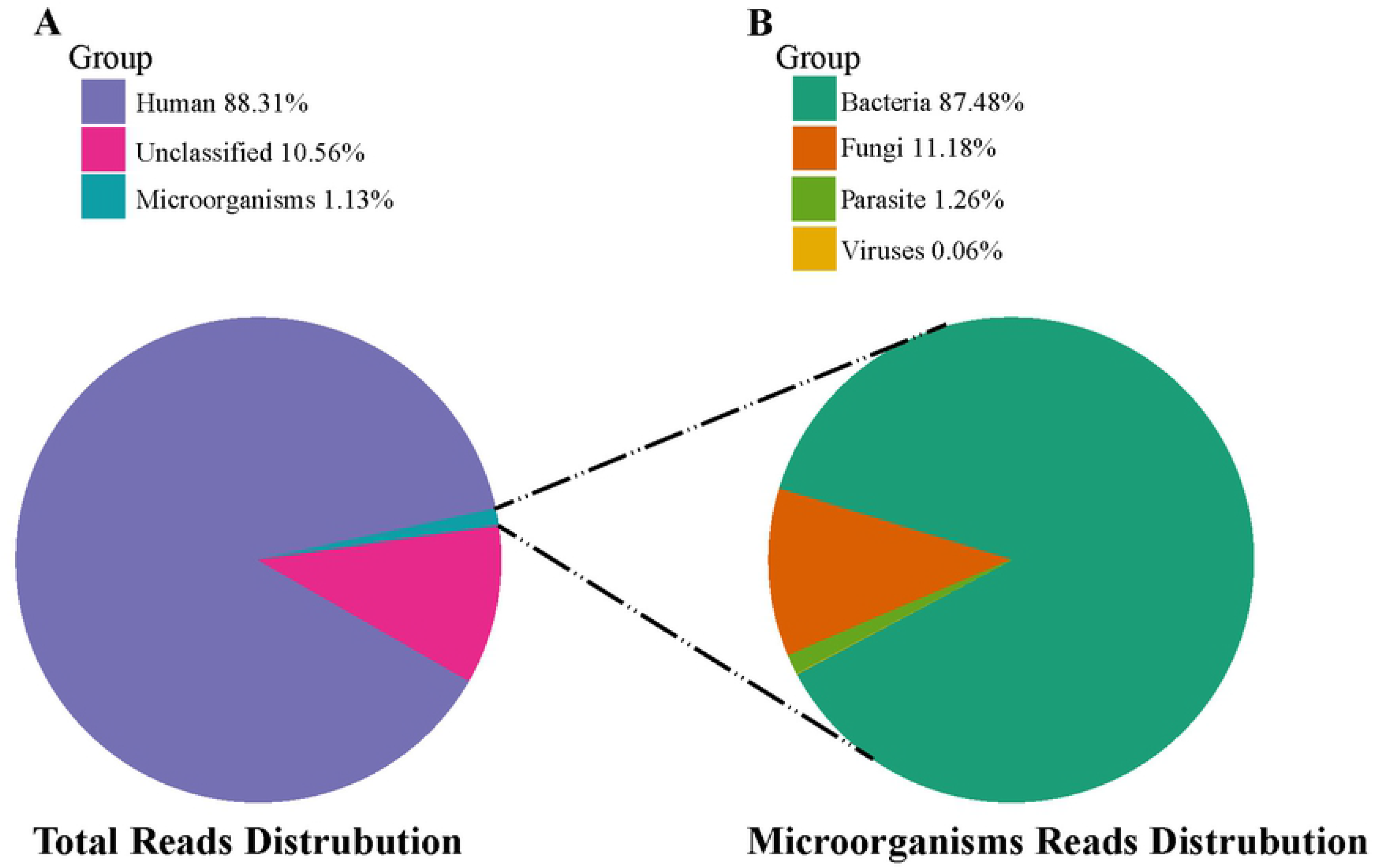
Description of mNGS sequencing results. (A) Reads distribution of total DNA in all environmental samples. (B) Distribution of pathogenic microorganisms reads in the absence of human and unclassified reads.

### Differences in Flora Distribution and Principal Component Analysis

This study analyzed the differences in the distribution of different types of DNA on instrument for joint arthroplasty in different medical institutions. We use PCA (Principal Component Analysis) method to extract common features from the original data and construct principal component factors, which reflect the differences and consistency of bacterial species between medical institutions in different regions. The results showed that there were three obvious clusters, indicating that the distribution of bacteria on the surface of instruments was different among the 7 medical institutions (Fig 2).

**Fig 2.**
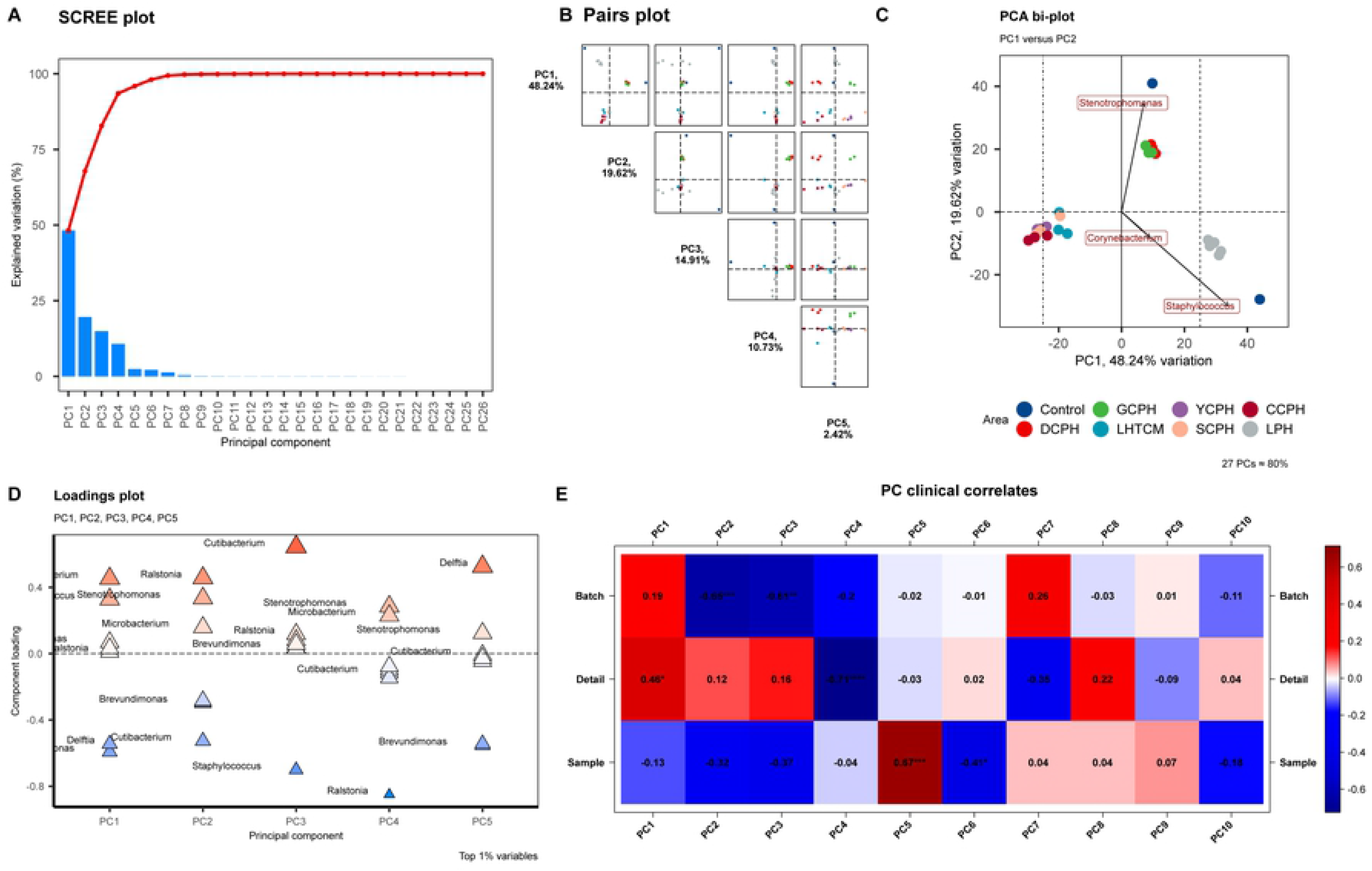
Principal Component Analysis of the microbiome composition. (A) Scree plot, determine the number of principal components to keep in a principal component analysis. (B) Pairs plot demonstrates the dispersion and aggregation of samples at different PC scales. (C) PCA bi plot, demonstrates the dispersion and aggregation of samples at two main PC scales. (D) Loading plot, determine the variables that drive variation among each PC. (E) PC clinical correlates, Correlate the principal components back to the clinical data (Batch, Detail, Sample).

To further clarify the differences in the distribution of bacteria on the surface of orthopedic instruments in different hospitals and the characteristic flora of each medical institution, we used a Venn diagram to show the common and unique types of bacteria among different medical institutions (Fig 3A). The results showed that there were 465 species of bacteria in all regions, and Liaocheng People’s Hospital had a maximum of 340 species of bacteria, followed by County People’s Hospital with 169 species. Afterward, we exhibited the differences in bacterial distribution among medical institutions through clustering heatmaps. From Fig 3B, we found a similar distribution of microbiota in the three samples from different instruments in the same hospital, and the differences between hospitals are very noticeable. There was no significant difference between LPH 1-3 (originally sampled from Liaocheng People’s Hospital) and LPH 4-6 (added human cells). Three genera of “*Cutibacterium*”, “*Acinetobacter*” and “*Staphylococcus*” are commonly found in the sequencing data, which are also universal background genera in routine sequencing. After that, we analyzed the main bacterial genera whose sequences accounted for more than 1% in each region and drew the stacked histogram of the genera. The results can be seen that the distribution of the main environmental genera in different regions was significantly different. The distribution of predominant genera differed substantially between hospitals (Fig 3C). The complete sequencing raw data are shown in S1 Tables. For the comparison chart of fungi, viruses, and parasites, see S1-S6 Figs.

**Fig 3.**
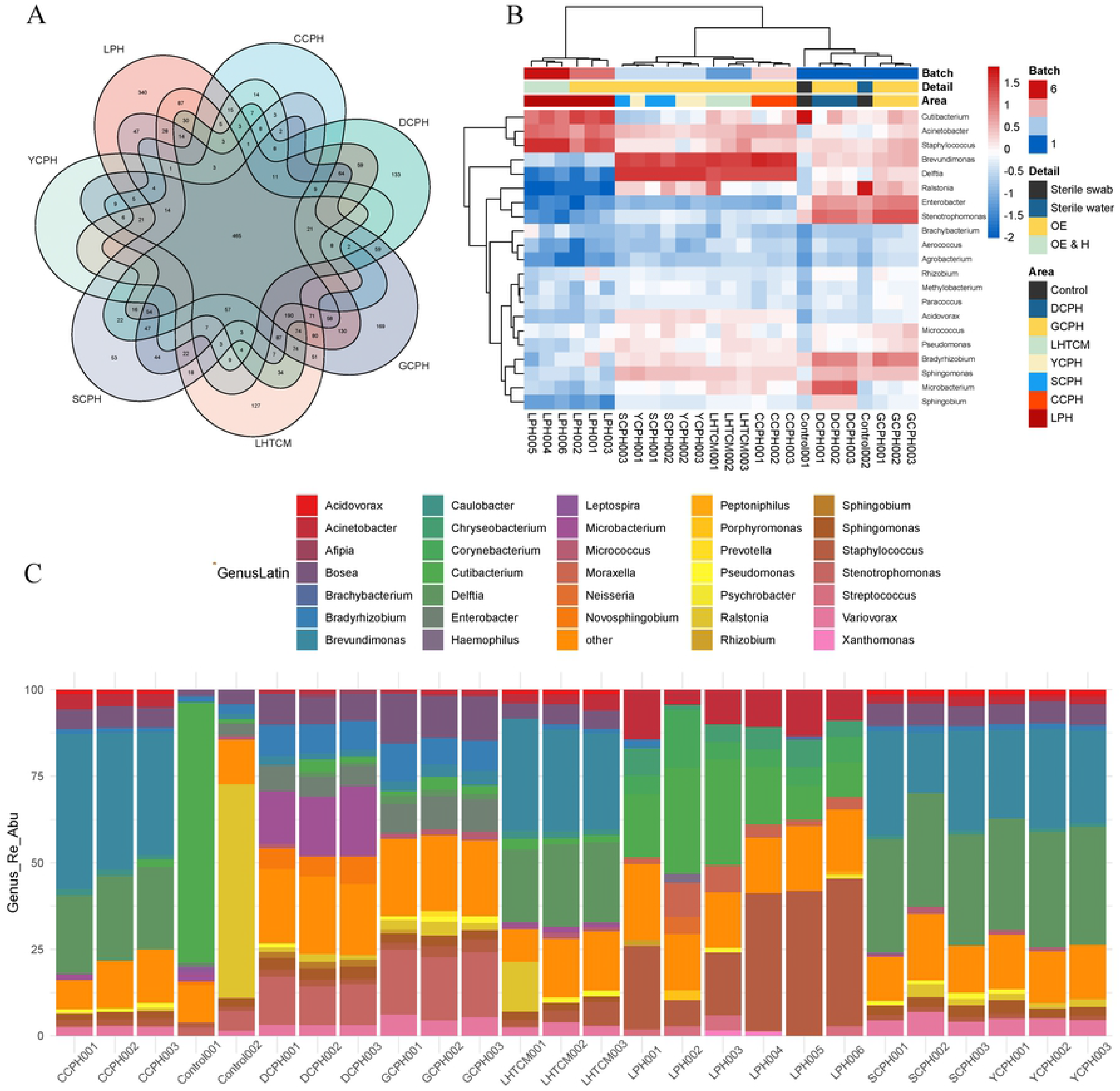
The distribution of bacteria on the surfaces of instruments varied between hospitals, as did the characteristic flora of each healthcare facility. (A) Venn diagram, show the common and unique types of bacteria among different medical institutions. (B) Heat map of the relative abundance at the genus level of bacteria, Demonstrates similarities and differences in the distribution of bacterial genus levels in different medical institutions. (C) Cumulative map of the relative abundance of bacterial spp. levels in different healthcare facilities, demonstrates the distribution of the dominant genera in different healthcare facilities.

## Discussion

At present, although a small number of studies have focused on the contamination of background nucleic acid fragments in mNGS sequencing, most of them are limited to empirical speculation <u>[15]</u>. To our knowledge, this study is the first to apply a data-based approach to study the impact of nucleic acid fragments in the instrument environment during mNGS on the diagnosis of PJI. Although we used primers that can amplify pathogen DNA fragments in the mNGS sequencing process, human-derived sequence fragments still accounted for the vast majority, which is consistent with the sequencing results of general joint samples. The main environmental pollution DNA fragments found in this study were bacterial fragments, while viral, fungal, and parasitic fragments were extremely rare. This is consistent with previous research findings and general clinical experience that most PJIs are due to bacterial infections <u>[15]</u>. There were no obvious differences in the sequences of the detected fungi and viruses, which may be related to their fewer fragments. The detected residues are considered to be more likely to be mismatched bases. Furthermore, we found significant differences in the distribution of bacterial DNA fragments on the surfaces of instruments in different hospitals, which is likely to be related to bacteria from previous PJI cases handled by the device. At present, conventional disinfection methods can effectively remove all active pathogens, and no pathogens were cultured in this study. Nevertheless, DNA fragments are difficult to remove by conventional sterilization means due to their inherent stability. Therefore, DNA fragments on the device will be stored for a long time and may affect the mNGS results of subsequent PJI patients.

In a recent study on the etiology of PJI, the results of Benito *et al.* identified *Staphylococcus* as the most common cause of infection <u>[16]</u>. In addition, other gram-positive cocci (8-9%), gram-negative bacilli (6%), anaerobic bacteria (4%) and Candida (1%) may also cause PJI <u>[16, 17]</u>. Rare microorganisms that cause PJI also include, but are not limited to, *Aspergillus fumigatus*, *Actinomyces*, and *Mycoplasma hominis* <u>[18, 19]</u>. Considering the complexity of PJI infection, the application of mNGS technology to detect pathogens has positive significance for the symptomatic treatment and prognosis of PJI. The high sensitivity of mNGS leads to the possibility of false positives caused by any nucleic acid residues of pathogenic microorganisms exposed during the sampling process. Thus, it is necessary to introduce the concept of BML and monitor their possible sources. During the sampling process, the containers used are sterile and nucleic acid-free according to standard procedures. Generally, no nucleic acid residues of pathogenic microorganisms are introduced. To exclude the influence of nucleic acid residues in the detection reagents, we set up two synchronized negative controls, including an unsampled swab control (Control 1) and pure water (Control 2). The results showed that the detection reagent did not affect the distribution of microorganisms among different hospitals. Therefore, medical institutions must establish a localized mNGS BML and the rules for distinguishing it from normal pathogenic bacteria. Besides, the detection, assessment, and periodic testing of nucleic acid fragments in reagents can help minimize cross-contamination between tested samples <u>[2, 12, 20]</u>.

The current mainstream mNGS report interpretation methods tend to be comprehensively analyzed by a team composed of clinicians, laboratory technicians, and bioinformatics personnel. Even so, the fragment contamination of environmental nucleic acid cannot be well differentiated, resulting in false positives <u>[21]</u>. Most of the interfering fragments found in this study are pathogens that are not common in clinical infections (eg *Corynebacterium*, *Novosphingobium*, *Sphingomonas*), which are relatively easier to distinguish from PJI-infected bacteria. However, it cannot be ruled out that in immunocompromised patients, the joints will be infected with these special pathogenic bacteria. On the other hand, we also encountered residues of nucleic acid fragments with strong interference (eg, *Staphylococcus*, *Acinetobacter*), which overlapped with common pathogens of PJI. These fragments make the interpretation of the results more complicated, and better methods need to be explored to judge or exclude. One study summarized common background bacterial species through experience, including *Acinetobacter*, *Streptococcus*, *Propionibacterium*, *Bradyrhizobium*, *Dolosigranulum*, etc. <u>[15]</u>. However, this is only observational and has not been statistically and verified by data. The distribution of genus was significantly different from our study. It can be seen that the background bacteria of different platforms and different regions are not completely consistent.

Through the establishment of BML of different medical institutions, we can provide a strong reference for the reporting and interpretation of clinical specimens. However, it should be noted that some contaminating bacterial groups may be indistinguishable from infectious pathogens even by establishing a library. Identifying and removing contaminant DNA requires a complex set of methodologies. Reliable bioinformatics tools and databases are needed to decide whether detected microorganisms are the cause of infection or human-derived sequence contamination as well as background contamination. In addition, we tested samples from the same source with and without the introduction of human sequences and confirmed no significant differences. This may be related to the high concentration of human DNA on the surface of the device sample itself. Analyzing bioinformatic readouts (such as coverage, abundance, etc.) will play a key role in the adoption and utility of this method. However, in low-level infections caused by low-abundance pathogens, the presence of contaminants can make the correct interpretation of mNGS readings difficult. Furthermore, we run into the problem of increasing the amount of off-camera data to increase sensitivity, but the increased sensitivity of mNGS confounds the boundaries we draw in terms of organisms or relatedness of identified organisms. In special cases, factors such as the patient’s condition, the joint to be replaced, the different geographic regions, and the time from surgery to consultation should be taken into account <u>[22]</u>. The pathogens themselves should also be characterized, for example, coagulase-negative *Staphylococci* species are part of the human skin microbiome and include a large group of bacteria such as *Staphylococcus epidermidis* and *Staphylococcus ludens* <u>[23]</u>. Another example is *Dermatobacter acnes*, a common skin-colonizing bacterium that can cause joint infections. Even with bacterial load thresholds set for these specific pathogens, it is difficult to differentiate between true pathogens and contaminants. Therefore, we need a comprehensive analysis combining multiple factors, such as a higher proportion of prosthetic shoulder joint infections caused by *D. acnes* compared to other joints <u>[24]</u>.

The study still has several limitations. That is, we have collected samples from different hospitals in the same period, but have not conducted continuous monitoring on the background bacteria of the samples in different periods. Similar to the regular monitoring of environmental microorganisms in medical institutions based on routine bacterial culture, periodic sequencing of residual nucleic acids from instruments for joint arthroplasty may become possible in the future. In addition, due to the low sequence of residual fragments of viruses and fungi, false-positive interference informal testing cannot be completely ruled out. Among them, it is difficult for fungi to extract gene fragments due to their thick cell wall, especially filamentous fungi. Judgment rules cannot be treated equally with gene sequence fragments of bacteria. In other words, the diagnostic thresholds of these microbial infections require the establishment of more refined rules and the accumulation of experience to distinguish the correspondence between PJI or false positives and clinical decisions.

## Conclusion

The surface of the instrument for joint replacement sterilized by the routine process does not contain any active microorganisms, but there are residual microbial nucleic acid fragments, which will interfere with the mNGS test results. The residual microbial nucleic acid fragments on the surface of joint replacement devices are mainly bacterial DNA, and there are differences in the distribution of bacteria between different medical institutions, which may be related to the historical cases handled by the medical treatment. In the current form, the establishment of independent BML of different medical institutions can effectively improve the accuracy of mNGS diagnosis and help to exclude the interference of background microorganisms.

## Supporting information

**S1Table. Details of metagenomics analysis results.** Listed for each sample are the Species_Name, Coverage, CovRate (coverage rate), Depth, Species_SMRN (stringently mapped read numbers to species), Genus_Name, Genus_SMRN (stringently mapped read numbers to genus), Species_Re_Abu (relative abundance of species), Genus_Re_Abu (relative abundance of genus), Species_Abs_Abu (absolute abundance of species), Genus_Abs_Abu (absolute abundance of genus), Shannon_index (diversity index).

Supple

**S1 Fig. Principal component analysis of the distribution of fungi communities.** (A) Scree plot, determine the number of principal components to keep in a principal component analysis. (B) Pairs plot demonstrates the dispersion and aggregation of samples at different PC scales. (C) PCA bi plot, demonstrates the dispersion and aggregation of samples at two main PC scales. (D) Loading plot, determine the variables that drive variation among each PC. (E) PC clinical correlates, Correlate the principal components back to the clinical data (Batch, Detail, Sample).

**S2 Fig. Heat map and cumulative map of the relative abundance of fungi.** Heat map of the relative abundance at the genus level of fungi, Demonstrates similarities and differences in the distribution of genus levels in different medical institutions. Cumulative map of the relative abundance of fungi spp. levels in different healthcare facilities, demonstrates the distribution of the dominant genera in different healthcare facilities.

**S3 Fig. Principal component analysis of the distribution of virus communities.** (A) Scree plot, determine the number of principal components to keep in a principal component analysis. (B) Pairs plot demonstrates the dispersion and aggregation of samples at different PC scales. (C) PCA bi plot, demonstrates the dispersion and aggregation of samples at two main PC scales. (D) Loading plot, determine the variables that drive variation among each PC. (E) PC clinical correlates, Correlate the principal components back to the clinical data (Batch, Detail, Sample).

**S4 Fig. Heat map and cumulative map of the relative abundance of viruses.** Heat map of the relative abundance at the species level of viruses, Demonstrates similarities and differences in the distribution of species levels in different medical institutions. Cumulative map of the relative abundance of virus species levels in different healthcare facilities, demonstrates the distribution of the dominant virus in different healthcare facilities.

**S5 Fig. Principal component analysis of the distribution of parasite communities.** (A) Scree plot, determine the number of principal components to keep in a principal component analysis. (B) Pairs plot demonstrates the dispersion and aggregation of samples at different PC scales. (C) PCA bi plot, demonstrates the dispersion and aggregation of samples at two main PC scales. (D) Loading plot, determine the variables that drive variation among each PC. (E) PC clinical correlates, Correlate the principal components back to the clinical data (Batch, Detail, Sample).

**S6 Fig. Heat map and cumulative map of the relative abundance of parasites.** Heat map of the relative abundance at the species level of parasites, demonstrates similarities and differences in the distribution of species levels in different medical institutions. Cumulative map of the relative abundance of parasite species levels in different healthcare facilities, demonstrates the distribution of the dominant parasites in different healthcare facilities.

## Notes

### Competing Interest Statement

The authors have declared no competing interest.

